# A global genomic approach uncovers novel components for twitching motility-mediated biofilm expansion in *Pseudomonas aeruginosa*

**DOI:** 10.1101/372805

**Authors:** Laura M. Nolan, Cynthia B. Whitchurch, Lars Barquist, Marilyn Katrib, Christine J. Boinett, Matthew Mayho, David Goulding, Ian G. Charles, Alain Filloux, Julian Parkhill, Amy K. Cain

## Abstract

*Pseudomonas aeruginosa* is an extremely successful pathogen able to cause both acute and chronic infections in a range of hosts, utilizing a diverse arsenal of cell-associated and secreted virulence factors. A major cell-associated virulence factor, the Type IV pilus (T4P), is required for epithelial cell adherence and mediates a form of surface translocation termed twitching motility, which is necessary to establish a mature biofilm and actively expand these biofilms. *P. aeruginosa* twitching motility-mediated biofilm expansion is a coordinated, multicellular behaviour, allowing cells to rapidly colonize surfaces, including implanted medical devices. Although at least 44 proteins are known to be involved in the biogenesis, assembly and regulation of the T4P, with additional regulatory components and pathways implicated, it is unclear how these components and pathways interact to control these processes. In the current study, we used a global genomics-based random-mutagenesis technique, <u>tra</u>nsposon <u>d</u>irected insertion-site <u>s</u>equencing (TraDIS), coupled with a physical segregation approach, to identify all genes implicated in twitching motility-mediated biofilm expansion in *P. aeruginosa*. Our approach allowed identification of both known and novel genes, providing new insight into the complex molecular network that regulates this process in *P. aeruginosa*. Additionally, our data suggests a differential effect on twitching motility by flagellum components based upon their cellular location. Overall the success of our TraDIS approach supports the use of this global genomic technique for investigating virulence genes in bacterial pathogens.

## Introduction

*P. aeruginosa* is a leading cause of health-care associated infections and is the major cause of mortality in patients with cystic fibrosis (CF) (1). This bacterium’s success as a pathogen is mainly attributed to its ability to produce a plethora of cell-associated and secreted virulence factors (2). Type IV pili (T4P) are major cell-associated virulence factors of *P. aeruginosa*, promoting both attachment to host epithelial cells and a form of flagella-independent surface translocation, termed twitching motility (3). Twitching motility is a complex and co-ordinated multicellular phenomenon, which in *P. aeruginosa*, results in active biofilm expansion (4, 5). This active expansion can result in the spread of infection within host tissues and along implanted medical devices (6, 7).

The biogenesis, assembly and regulation of the T4P for mediating twitching motility-mediated biofilm expansion requires at least 44 different proteins (3, 4, 8). The components involved in biogenesis and assembly of T4P are encoded by *pilA*, *B*, *C*, *D*, *E*, *F*, *M*, *N*, *O*, *P*, *Q*, *T*, *U*, *V*, *W*, *X*, *Y1*, *Y2*, *Z*, and *fimT*, *U*, *V*. The T4P is composed of multiple PilA monomers, which are assembled by the T4P biogenesis machinery. This machinery is composed of the motor, alignment and pilus subcomplexes (3). At the inner membrane, the motor subcomplex is composed of a platform protein PilC and the cytoplasmic ATPases PilB and PilT, which are responsible for pilus elongation and retraction, respectively (9, 10). The alignment subcomplex composed of PilM, PilN and PilO forms a connection between the motor subcomplex and the outer membrane associated secretin complex of PilP and PilQ (11-14). Pilus extension is mediated by the combined activity of PilZ, FimX, and the ATPase PilB, while the ATPases PilT and/or PilU are responsible for pilus retraction (15-21).

Expression of these T4P genes is controlled by several systems in *P. aeruginosa* including the two-component sensor-regulator pairs, PilS/PilR (22, 23) and FimS/AlgR (24). The highly complex Chp chemosensory system encoded by the *pilGHIJK-chpABC* gene cluster (25-28) is involved in regulating the motors which control T4P extension and retraction in response to environmental signals (29), and is related to the Che chemosensory signal transduction system in *E. coli* which regulates flagella-mediated chemotaxis (30). Other regulatory components include: the virulence factor regulator Vfr (homologous to the *E. coli* catabolic repressor protein (CRP)) (31, 32); FimL which appears to intersect with the Chp chemosensory system and Vfr regulatory cascade (33); Crc, the catabolic repressor control protein (34); FimX (21, 35); FimV (36); as well as PocA, PocB and TonB3 (37, 38). Additionally, the regulation of T4P biogenesis, assembly and twitching motility-mediated biofilm expansion is further complicated by the contribution of the small intracellular signalling molecules 3’,5’-cyclic adenosine monophosphate (cAMP) and 3’,5’-cyclic diguanylic acid (c-di-GMP) (35, 39, 40). Clearly the regulation of T4P biogenesis and assembly, and twitching motility-mediated biofilm expansion is complex, and while multiple components and signalling cascades have been implicated, it remains unclear precisely how these components and pathways intersect. Therefore, we predict that other, currently uncharacterised proteins and pathways provide the links between these known components and pathways that regulate T4P biogenesis, assembly and twitching motility.

Here we adapted a global genomics-based approach termed <u>tra</u>nsposon <u>d</u>irected insertion-site <u>s</u>equencing (TraDIS) (41, 42) to identify all genes involved in twitching motility-mediated biofilm expansion in *P. aeruginosa*. TraDIS is a powerful method which utilises high-throughput sequencing of dense random transposon mutant libraries to identify all genes involved in any selective condition (42, 43). TraDIS (and related transposon-insertion sequencing methods) has proved successful for assaying a number of phenotypes in *P. aeruginosa*, including antibiotic resistance (44), stress conditions (45) and *in vivo* wound infection (46). However, this is the first time TraDIS has been used to investigate biofilm expansion in *P. aeruginosa.* The success of our systematic and global approach for investigating this aspect of *P. aeruginosa* virulence demonstrates that TraDIS-based separation is a powerful method to generate a comprehensive catalogue of genes involved in motility and pathogenesis-associated phenotypes, and the physical segregation approach used here can be applied more broadly to study bacterial phenotypes other than simple survival in selective conditions.

## Materials and Methods

### Bacterial strains, media and twitching motility assays

Strains used in this study were *P. aeruginosa* strain PA14, mutants from the PA14 non-redundant transposon mutant library (47) (see Table S2) and *E. coli* S17-1 containing the mini-Tn*5*-pro plasmid (48). *P. aeruginosa* strain PAK (Filloux lab collection) was used for generation of mutant strain PAK_05353 (orthologue of PA5037/PA14_66580) generated by allelic exchange mutagenesis as described previously (49, 50). Additionally PA14*pilR*::mar2xT7 (47) was used as a pilin negative control for TEM analysis, and PAKΔ*pilQ* (gifted by Stephen Lory) and PAK*pilA*:TcR (51) as negative controls for Western blot analyses.

*P. aeruginosa* and *E. coli* were cultured on Luria–Bertani (LB) (52) broth solidified with agar at 1.5 % or 1 % (for twitching motility sub-surface assays) and grown overnight at 37 °C. Cultures were grown in either cation-adjusted Mueller Hinton broth (CAMHB) or LB broth, and incubated overnight at 37 °C, with shaking at 250 rpm. Antibiotic concentrations used were gentamicin 15 μg/mL for plasmid maintenance in *E. coli* and for recovery of PA14 transposon mutants from library glycerol stocks; after initial recovery of PA14 transposon mutants from glycerol no antibiotic selection was used. Twitching motility-mediated biofilm expansion was assayed in sub-surface stab assays as described previously (40). Generated TraDIS library transposon mutants were recovered on 1x Vogel-Bonner Media (VBM) (a 10x solution contains (MgSO4.7H2O (8 mM), citric acid (anhydrous) (9.6 mM), K_2_HPO_4_ (1.7 mM), NaNH_5_PO_4._4H20 (22.7 mM), pH 7, and filter sterilized) with 1.5 % agar containing gentamycin at 100 μg/mL.

### Planktonic growth assays

Planktonic growth of *P. aeruginosa* was followed by recording changes in OD_600nm_ for 20 h, with incubation at 37 °C and shaking at 250 rpm. Cells were grown in 96-well microtitre plates with LB to replicate the conditions in the sub-surface twitching motility assays, or minimal media (1x M63 ((NH_4_)_2_SO_4_ (15 mM); KH_2_PO_4_ (22 mM); and K_2_HPO_4_ (40 mM)) supplemented with MgSO_4_ (1 mM), casamino acids (0.05%) and glucose (0.4%)) to replicate the conditions in the attachment assays.

### Submerged biofilm assays

Overnight cultures were diluted to OD_600nm_ = 0.1 into microtitre plates with 1x M63 media plus supplements (as for minimal media growth assays above) and incubated statically at 37 °C for 18 h. Planktonic growth was then removed and the remaining attached cells stained with crystal violet (10% (v/v)) for at least 10 min, statically, at room temperature. Unbound crystal violet stain was removed and the plate washed twice prior to extraction of crystal violet dye with ethanol (95 %). OD_600nm_ of the crystal violet dye was then used to quantify the levels of attached cells.

### DNA manipulation

DNA isolation was performed using the PureLink Genomic DNA mini kit (Life Technologies) except for TraDIS library genomic DNA isolation (see below). Isolation of plasmid DNA was carried out using the QIAprep spin miniprep kit (Qiagen). Primers (Sigma) used are shown in Table 3. DNA fragments were amplified with either KOD Hot Start DNA Polymerase (Novagen) or standard Taq polymerase (NEB) as described by the manufacturer with the inclusion of Betaine (Sigma) or DMSO (Sigma). Restriction endonucleases were used according to the manufacturer’s specifications (Roche). DNA sequencing was performed by GATC Biotech.

**Table 1.** TraDIS identifies genes known to be involved twitching motility.

| Gene name | PA14 orthologue | Gene product | TraDIS log-fold change* |
| --- | --- | --- | --- |
| algR | PA14_69470 | alginate biosynthesis regulatory protein AlgR | -10.21 |
| chpA | PA14_05390 | ChpA | -12.74 |
| chpB | PA14_05400 | ChpB | +7.82 |
| chpC | PA14_05410 | putative chemotaxis protein methyltransferase | -1.08<sup>#</sup> |
| crc | PA14_70390 | catabolite repression control protein | -12.78 |
| fabF1 | PA14_25690 | beta-ketoacyl-acyl carrier protein synthase II | -3.82 |
| fimL | PA14_40960 | pilin biosynthetic protein | -10.25 |
| algZ | PA14_69480 | alginate biosynthesis protein AlgZ/FimS | -6.79 |
| fimT | PA14_60270.1 | type 4 fimbrial biogenesis protein FimT | -1.24<sup>#</sup> |
| fimU | PA14_60280 | type 4 fimbrial biogenesis protein FimU | -12.53 |
| fimV | PA14_20860 | putative T4P pilus assembly protein FimV | +1.04 |
| fimX | PA14_65540 | conserved hypothetical protein | -6.89 |
| pocB | PA14_25500 | conserved hypothetical protein | NA |
| pilA | PA14_58730 | type IV pilin structural subunit | +0.86<sup>#</sup> |
| pilB | PA14_58750 | type 4 fimbrial biogenesis protein PilB | -8.85 |
| pilC | PA14_58760 | type 4 fimbrial biogenesis protein pilC | -10.62 |
| pilD | PA14_58770 | type 4 prepilin peptidase PilD | -6.46 |
| pilE | PA14_60320 | type 4 fimbrial biogenesis protein PilE | -9.70 |
| pilF | PA14_14850 | type 4 fimbrial biogenesis protein PilF | -5.29 |
| pilG | PA14_05320 | type IV pili response regulator PilG | -8.64 |
| pilH | PA14_05330 | type IV pilus response regulator PilH | -3.17 |
| pilI | PA14_05340 | type IV pili signal transduction protein PilI | -10.18 |
| pilJ | PA14_05360 | type IV pili methyl-accepting chemotaxis protein | -7.73 |
| pilK | PA14_05380 | methyltransferase PilK | -0.18<sup>#</sup> |
| pilM | PA14_66660 | type 4 fimbrial biogenesis protein PilM | -6.22 |
| pilN | PA14_66650 | type 4 fimbrial biogenesis protein PilN | -6.86 |
| pilO | PA14_66640 | type 4 fimbrial biogenesis protein PilO | -11.35 |
| pilP | PA14_66630 | type 4 fimbrial biogenesis protein PilP | -6.96 |
| pilQ | PA14_66620 | type 4 fimbrial biogenesis OM protein PilQ | -6.20 |
| pilR | PA14_60260 | two-component response regulator PilR | -12.27 |
| pilS | PA14_60250 | kinase sensor protein of two component regulatory | -6.48 |
| pilT | PA14_05180 | twitching motility protein PilT | -9.69 |
| pilU | PA14_05190 | twitching motility protein PilU | -8.54 |
| pilV | PA14_60280.1 | type 4 fimbrial biogenesis protein PilV | -13.14 |
| pilW | PA14_60290 | type 4 fimbrial biogenesis protein PilW | -8.71 |
| pilX | PA14_60300 | type 4 fimbrial biogenesis protein PilX | -8.86 |
| pilY1 | PA14_60310 | type 4 fimbrial biogenesis protein PilY1 | -7.56 |
| pilY2 | PA14_60310.1 | type 4 fimbrial biogenesis protein PilY2 | -9.50 |
| pilZ | PA14_25770 | type 4 fimbrial biogenesis protein PilZ | -9.03 |
| ppk | PA14_69230 | polyphosphate kinase | +3.94 |
| rpoN | PA14_57940 | RNA polymerase sigma-54 factor | NA |
| rpoS | PA14_17480 | sigma factor RpoS | -5.85 |
| tonB3 | PA14_05300 | TonB3 | -7.53 |
| vfr | PA14_08370 | cyclic AMP receptor-like protein | -9.30 |

\*A positive log-fold change indicates that the gene product has a negative effect on twitching motility, while a negative log-fold change has a positive effect on twitching motility; # result is not significant (Q-value is >0.05) but confirmed by visual inspection to have differential insertion levels during twitching motility-mediated biofilm expansion; NA – not assayed due to a minimal insertion density in these genes in our starting base library.

**Table 2.** Transposon mutants with increased or decreased TM compared to PA14 wildtype for further analysis.

| PA14 transposon mutant* | PAO1 orthologue | Functional class<sup>#</sup> | Description |
| --- | --- | --- | --- |
| PAMr_nr_mas_07_1:C3 | *prlC* | Amino acid transport and metabolism | Zn-dependent oligopeptidase |
| PAMr_nr_mas_04_1:A9 | *lon* | Posttranslational modification, protein turnover, chaperones | ATP-dependent Lon protease |
| PAMr_nr_mas_07_1:D4 | *kinB* | Signal transduction mechanisms | Signal transduction histidine kinase |
| PAMr_nr_mas_12_4:D5 | *cyaB* | Signal transduction mechanisms | Adenylate cyclase |
| PAMr_nr_mas_12_1:C6 | *pfpI* | Bacterial-type flagellar swarming motility; cellular response to antibiotic; single-species biofilm formation | Intracellular protease |
| PAMr_nr_mas_05_1:H3 | *fliG* | Cell motility | Flagellar motor switch protein |
| PAMr_nr_mas_09_4:G3 | *fliF* | Cell motility, intracellular trafficking, secretion, and vesicular transport | Flagellar basal body M-ring protein |
| PAMr_nr_mas_07_1:C7 | *motB* | Bacterial-type flagellar cell motility | Flagellar motor protein |
| PAMr_nr_mas_10_2:F2 | *motY* | Cell wall/membrane/envelope biogenesis | Sodium-dependent flagellar system protein |
| PAMr_nr_mas_09_3:C3 | *algU* | Sigma factor activity; negative regulation of bacterial-type flagellar cell motility; regulation of polysaccharide biosynthetic process | Sigma factor |
| PAMr_nr_mas_14_4:C8 | PA5037 (PA14_66580) | ATPase activity; Type II secretory pathway, component ExeA | Hypothetical |

\* Plate and well for mutants from PA14 transposon mutant collection (Liberati et al. (2006)); <sup>#</sup> Colours denote mutants which have similar predicted functional classes for the gene product using Clusters of Orthologous Groups (COGs).

**Table 3.**
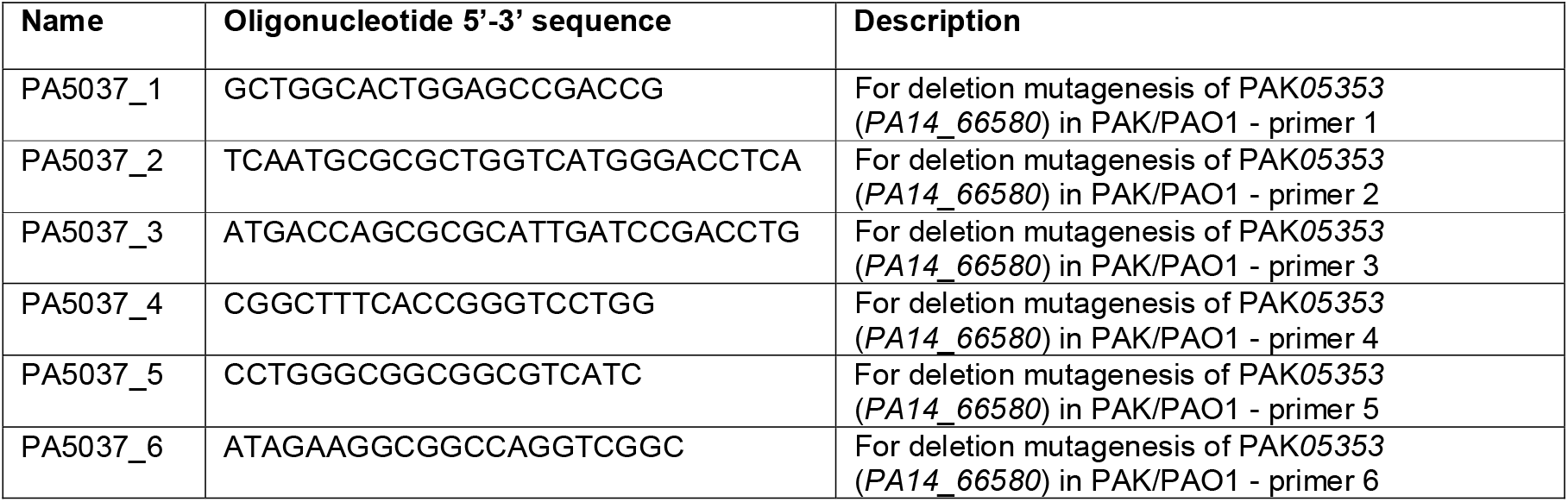
Oligonucleotides used in this study.

### Preparation of samples for PilQ immunoblotting

Preparation of whole cell samples for PilQ analysis were performed as described previously (28) with cells being harvested from plates grown for 20 h at 37 °C on LB agar. For analysis of PilQ multimerization samples were only boiled for 2 min at 95 °C in Laemmli loading buffer prior to loading on the SDS-PAGE gel. All other Western blot samples were boiled for 10 min at 95 °C in Laemmli loading buffer prior to loading on the SDS-PAGE gel

### Western Blot analysis

SDS-PAGE and western blotting were performed as described previously (50). Proteins were resolved in 8%, 10%, 12% or 15% gels using the Mini-PROTEAN system (Bio-Rad) and transferred to Nitrocellulose membrane (GE Healthcare) by electrophoresis. Membranes were blocked in 5% milk (Sigma) before incubation with primary antibodies. Membranes were washed with TBST (0.14 M NaCl, 0.03 M KCl and 0.01 M phosphate buffer plus Tween 20 (0.05% v/v)) before incubation with HRP-conjugated secondary antibodies (Sigma). The resolved proteins on the membrane blots were detected using the Novex ECL HRP Chemioluminescent substrate (Invitrogen) or the Luminata Forte Western HRP substrate (Millipore) using a Las3000 Fuji Imager. Membranes were probed with α-PilQ antibody (gifted by Stephen Lory), or α-RNAP antibody (Neoclone) and secondary anti-rabbit antibody for PilQ and anti-mouse antibody for RNAP.

### Transmission Electron Microscopy (TEM) assays

Log phase cultures (OD_600nm_ 0.25 to 0.5) were fixed in the planktonic state with 0.1 % glutaraldehyde and then spotted on a 400 mesh copper/palladium grid. Alternatively, cells were first spotted on a grid, incubated for 15 min at room temperature, and then fixed in a surface-associated state with 0.1 % glutaraldehyde. Preparations were then washed 3 times with water and negatively-stained twice with 1 % uranyl acetate. Images were taken with a FEI Morgagni 268(D) electron microscope.

### TraDIS library generation

A highly saturated transposon mutant library was generated in *P. aeruginosa* PA14 by large scale conjugation with an *E. coli* SM17-1 [mini-Tn*5*-pro]] donor which allowed for random insertion of a mariner transposon throughout the PA14 genome, and conferred gentamicin resistance in the recipient PA14 strain. The *E. coli* donor strain was grown in LB supplemented with gentamicin (15 μg/mL) overnight at 37 °C and the recipient PA14 strain was grown overnight at 37 °C in CAMHB. Equivalent amounts of both strains were spread uniformly on separate LB agar plates and incubated overnight at 37 °C for *E. coli* and at 43 °C under humid conditions for *P. aeruginosa*. The next day 1 *E. coli* donor plate was harvested and combined by extensive physical mixing on a fresh LB agar plate with 1 plate of harvested recipient PA14 strain. Conjugation between the two strains was achieved by incubation of the high-density mixture of both strains at 37 °C for 2 hours. The conjugation mix was then harvested, pelleted by centrifugation (10,000 *g*, 10 min, 4 °C), and resuspended in LB. The resuspended cells were recovered on 1x VBM agar supplemented with gentamicin (100 μg/mL) and incubated for 17 h at 37 °C. The numbers of mutants obtained were estimated by counting a representative number of colonies across multiple plates. Mutants on plates were recovered as a pool, resuspended in LB, pelleted by centrifugation (10,000 *g*, 10 min, 4 °C), and then resuspended in LB plus glycerol (15 % (v/v)) and stored at -80 °C. The protocol was repeated on a large scale until ~2 million mutants were obtained.

### TraDIS assay with mutant pool

The transposon mutant library pool was diluted 1:10 into 9 mL CAMHB in 10 separate 50 mL Falcon tubes which were covered with aeroseal to facilitate aeration within the culture and incubated at 37 °C overnight. Thirty mL LB agar (1.5 %) 90 mm plates were poured and allowed to set overnight at room temperature. The following morning the agar was flipped into a larger petri dish to expose the smooth underside set against the petri dish base which promotes rapid twitching motility-mediated biofilm expansion (53). 1.5 mL of overnight growth of the pooled transposon mutant library was pelleted by centrifugation (10,000 *g*, 3 min, 4 °C), and the whole pellet then spotted into the centre of the flipped agar plate. This was repeated for all 10 overnight cultures and performed in triplicate (i.e. a total of 30 plates). All plates were incubated under humid conditions at 37 °C for 65 h. To harvest mutants based upon their ability to undergo twitching motility-mediated biofilm expansion mutants were harvested from the inner, non-twitching zone and from the outer, active-twitching motility zone (see Figure S1A) for all 3 replicates. The cells from the inner and from the outer zones were harvested separately for all 3 replicates by resuspension in 5 mL LB, followed by centrifugation (10,000 *g*, 10 min, 4 °C), to pellet the cells. The supernatant was discarded and the cells used for genomic DNA extraction.

### Genomic DNA extraction for TraDIS library sequencing

Genomic DNA from the harvested pooled library pellets was resuspended in 1.2 mL lysis solution (Tris-HCl (10 mM), NaCl (400 mM) and Na_2_EDTA (2 mM), supplemented with Proteinase K in storage buffer (Tris-HCl (50 mM), glycerol (50 % (v/v)), NaCl (100 mM), EDTA (0.1 mM), CaCl_2_ (10mM), Triton X-100 (0.1% (w/v)) and DTT (1 mM)) to a concentration of 166 μg/ml. Cell lysis was achieved by incubation at 65 °C for 1 h, with occasional vortexing. The samples were then cooled to room temperature and RNA removed by addition of RNaseA (5 μg/ml) and incubation at 37 °C for 80 min. Samples were then placed on ice for 5 min. Each lysate was then split into 2 eppendorf tubes of ~600 μL per tube, and 500 μL NaCl (3 M) added to each tube. Cell debris were removed by centrifugation (10,000 *g*, 10 min, 4 °C) and 500 μL from each tube was added to 2 volumes of isopropanol to precipitate DNA. DNA was then collected by centrifugation (10,000 *g*, 10 min, 4 °C), with the pelleted DNA being washed twice in 70 % (v/v) ethanol. DNA was finally resuspended in 50 μl Tris-EDTA buffer.

### Generation of DNA sequencing libraries and library sequencing

TraDIS was performed using the method described in Barquist et al., (2016). The PCR primers used were designed in this study, for library construction (5’: AATGATACGGCGACCACCGAGATCTACACAGGTTGAACTGCCAACGACTACG and 3’: AATGATACGGCGACCACCGAGATCTACACAACTCTCTACTGTTTCTCCATACCCG) and sequencing TraDIS primers (5’: CGCTAGGCGGCCAGATCTGAT, and 3’: GGCTAGGCCGCGGCCGCACTTGTGTA) and during library amplification, plasmid block primers were used to prevent amplification of plasmid background (5’ ctagaagaagcttgggatccgtcgaccgatcccgtacacaagtagcgtcc–dideoxy and 3’ attccacaaattgttatccgctcacaattccacatgtggaattccacatgtgg-dideoxy). For the TM TraDIS sequencing, we used a MiSeq Illumina platform and 13.2 million 150 bp single-end sequencing reads were generated. Reads were mapped onto PA14 (accession number: CP000438) genome, and 10% of the 3’ end of each gene was discounted, and a 10 read minimum cut-off used to be included in the comparisons performed using EdgeR (54), using scripts from the Bio-TraDIS pipeline (Barquist et al., (2016); https://github.com/sanger-pathogens/Bio-Tradis). All sequences from the TraDIS assays are available in the European Nucleotide archive (ENA) under study accession number ERP001977 and individual ENA accessions of each sample are ERS427191-3 for the non-twitching cells, ERS427194-6 for the twitching cells and ERS427197-9 for the base library without selection.

### Downstream analysis of TraDIS results

KEGG enrichment analysis was performed in R. KEGG pathway annotations were retrieved using the KEGGREST package. A hypergeometric test was used to test for pathway enrichment in genes with higher (logFC > 4, q-value < 0.01) or lower (logFC < -4, q-value < 0.01) mutant abundance in the TraDIS assay.

## Results

### Confirmation of genes known to be involved in twitching motility

To identify genes involved in twitching motility-mediated biofilm expansion we generated a high-density random transposon mutant library in *P. aeruginosa* PA14 using conjugation of a Tn*5* minipro vector and gentamicin selection. We determined that this library consisted of 310,000 unique Tn*5* mutants by sequencing DNA from 10^9^ cells from the raw base library, in duplicate, without selection.

Approximately 10^9^ cells from an overnight culture of the pool of transposon mutants were concentrated and inoculated as a central spot on top of an inverted agar plate. These were incubated for 65 h at 37 °C under humid conditions to allow a twitching motility-mediated surface biofilm to form. An inverted agar plate was used to expose the smooth underside of the moist, set agar, which facilitates rapid twitching motility-mediated biofilm expansion and discourages other forms of motility (53). This colony biofilm assay was favoured over the subsurface twitching motility assay (53) as this assay allows a much greater number of cells to be recovered, thus allowing sufficient amounts of genomic DNA to be extracted for downstream sequencing. Transposon mutants were separated based upon their ability to expand via twitching motility, away from the site of inoculation, with cells being harvested from the inner, non-twitching section of the colony biofilm, and the outer, actively expanding edge (Figure S1A). The outer and inner zones from 10 plates were combined to form each replicate, and 3 replicates were performed over different days. Genomic DNA was extracted from both combined pools of mutants then separately sequenced to determine the number of insertions per gene, using a TraDIS approach, as described previously (42). The relative frequencies of transposon insertion in the non-twitching and twitching transposon mutant pools were compared as described previously (42) and using a cut off of log_2_FC=4 and a Q-value of <0.01 to identify genes with differential insertion levels during twitching motility-mediated biofilm expansion. This revealed 942 genes as having a putative role in twitching motility-mediated biofilm formation: 82 genes with increased insertions and 860 with decreased insertions (Table S1). 42 of the 44 genes known to be involved in twitching motility-mediated biofilm expansion were identified (3, 4, 8) (Table 1). The two genes which we could not assay in our TraDIS screen were *rpoN* (PA14_57940) and *pocB* (PA14_25500) due to a minimal insertion density in these genes in our starting base library.

### Identification of novel components involved in twitching motility-mediated biofilm expansion

From our TraDIS results we selected 39 genes that had not been previously implicated in twitching motility (Table S2) to phenotypically characterize using single transposon mutants from the non-redundant PA14 transposon mutant collection (47). We tested the ability of each mutant to undergo twitching motility using a sub-surface stab assay. For those target genes that had multiple transposon mutants available, we tested all mutants, bringing the total number of assayed mutants to 52 (Table S2). From these assays, we detected 32 transposon mutants which had significantly altered levels of twitching motility compared to wildtype (Figure S2).

We selected 11 transposon mutants to further characterize based on biological interest and especially dramatic changes in twitching ability (Table 2). The genes containing these transposon insertions appear to group into distinct functional classes including cellular metabolism, signal transduction, cytokinesis and flagella-mediated motility (Table 2). Biofilm and growth assays were conducted for these 11 selected mutants to determine whether there was any effect on submerged biofilm formation (Figure 1B) and also to determine if the observed twitching motility defect (Figure 1A) was due to a growth-related effect (Figure 1C). Of these only *kinB* was found to have decreased levels of biofilm formation compared to wildtype (Figure 1B), as reported previously (55). None of these transposon mutants had an altered ability to grow in the minimal medium used for the biofilm assay demonstrating that any alteration in biofilm formation was not a result of a growth-related effect (Figure S1B). Of these 11 transposon mutants *prlC, lon, kinB, fliF* and *fliG* had significant alterations in growth rate in LB media (the same media used for sub-surface twitching motility assays) compared to wildtype (Figure 1C). Specifically, *fliF*, *fliG* and *prlC* had a shorter lag time than wildtype, and *kinB* and *lon* had a longer lag time than wildtype, but all reached approximately the same final cell density (Figure 1C). Based upon these results a growth-related effect may account for some of the observed decrease in twitching motility for *kinB* and *lon*. However, the observed alterations in twitching motility for *prlC*, *PA14_66580, pfpI, fliG, motY* and *algU* are not accounted for by growth related defects. Of these AlgU has already been implicated in regulation of twitching motility via what appears to be an indirect mechanism (24). To our knowledge none of the remaining targets *prlC*, *PA14_66580, pfpI, fliG* and *motY* have been previously implicated in twitching motility in *P. aeruginosa*.

**Figure 1.**
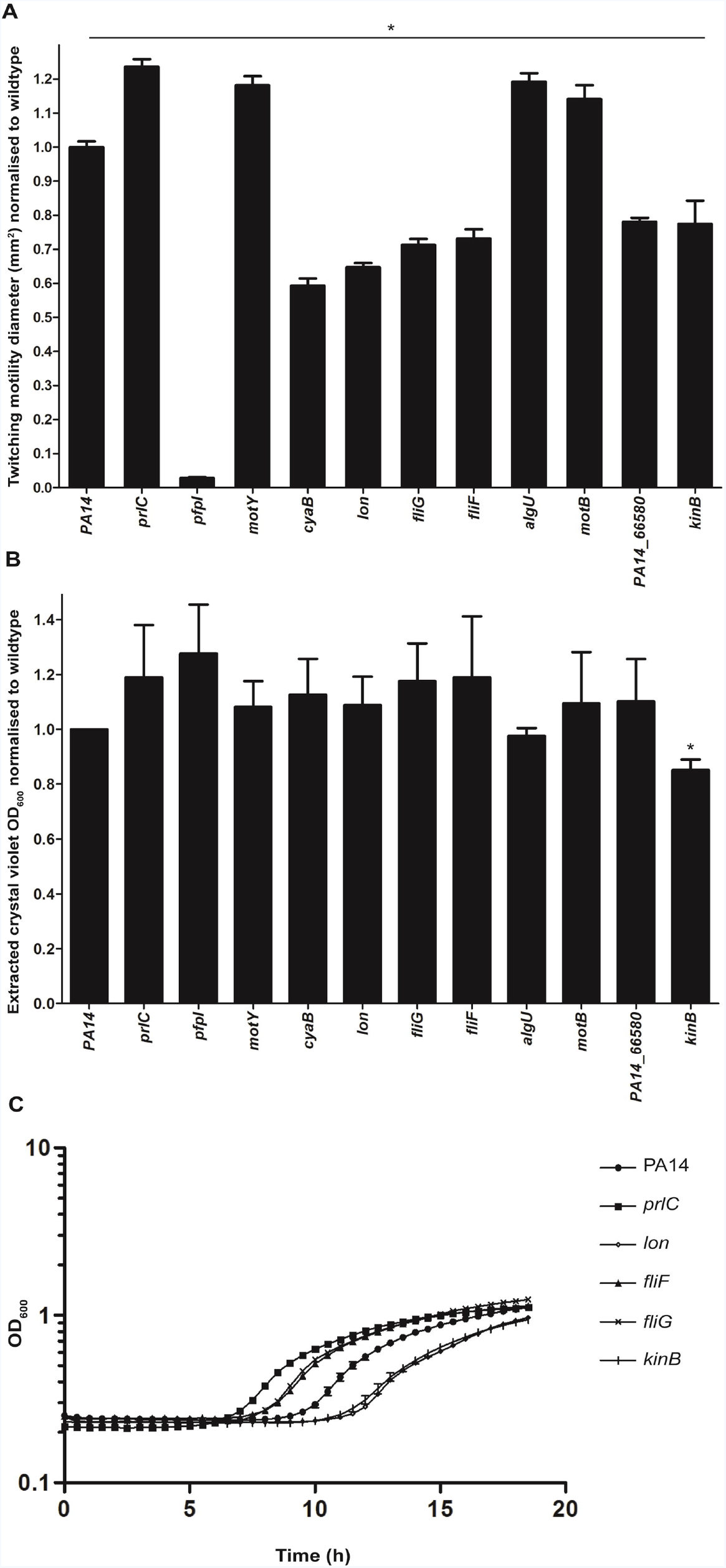
Characterization of twitching motility, biofilm and growth phenotypes for select transposon mutants. (A) Sub-surface twitching motility-mediated interstitial biofilm expansion at agar/plastic interface after 24 h incubation at 37 °C is presented as the mean surface area in mm^2^ ± standard error of the mean normalized against wildtype as obtained from 2 independent experiments performed in triplicate. Two-tailed student’s t-test, ^*^ (p<0.0005) compared to wildtype. (B) Submerged biofilm formation in 96-well microtitre plates after 18 h at 37 °C is presented as the mean OD_600nm_ reading ± standard error of the mean of extracted crystal violet staining normalized against wildtype, from 4 independent experiments performed in triplicate. Two-tailed student’s t-test, ^*^ (p<0.005) compared to wildtype. (C) Growth rates were determined by incubation of transposon mutants at 37 °C for 19 h in LB media. The growth rates for *prlC, lon, fliF, fliG* and *kinB* were significantly different (p<0.05) to wildtype (predicted by one-way ANOVA with Dunnett’s multiple comparison test). Levels of growth for *kinB, cyaB, pfpI, motB, motY, algU* and *PA14_66580* were not significantly different from PA14 wildtype (one-way ANOVA with Dunnett’s multiple comparison test).

### Visualizing the T4P of novel twitching motility gene targets using TEM

Alterations in twitching motility levels are commonly attributed to an increase or decrease in levels of expressed and/or assembled T4P and/or mislocalization of T4P. To determine whether the alterations in twitching motility of certain mutants result from abnormal levels or mislocalization of T4P, transmission electron microscopy (TEM) was used to visualize the pili. We investigated *prlC*, *PA14_66580, pfpI* and *motY*, as a flagellum-related representative, as well as PA14 wildtype and *pilR*, as respective positive and negative controls (Figure 2). These experiments revealed that PA14 wildtype, *PA14_66850, motY* and *prlC* possesses pili which were mainly polar (Figure 2A, C-G). Overall, the non-twitching *pfpI* mutant (Figure 1A) had no observable T4P (Figure 2B), both *PA14_66850* and *motY* had a reduced number of polar pili compared to wildtype (Figure 2C-D, F) and the hyper-twitching mutant *prlC* (Figure 1A) had extra-long pili, which in some cases appeared to intertwine with the observed flagella (Figure 2E) and an overall reduction in polar T4P levels compared to wildtype (Figure 2F). While PA14 wildtype, *PA14_66850* and *prlC* were found to possess non-polar T4P in a few cases, there was no difference in the numbers of non-polar T4P in the mutant strains compared to wildtype (Figure 2G).

**Figure 2.**
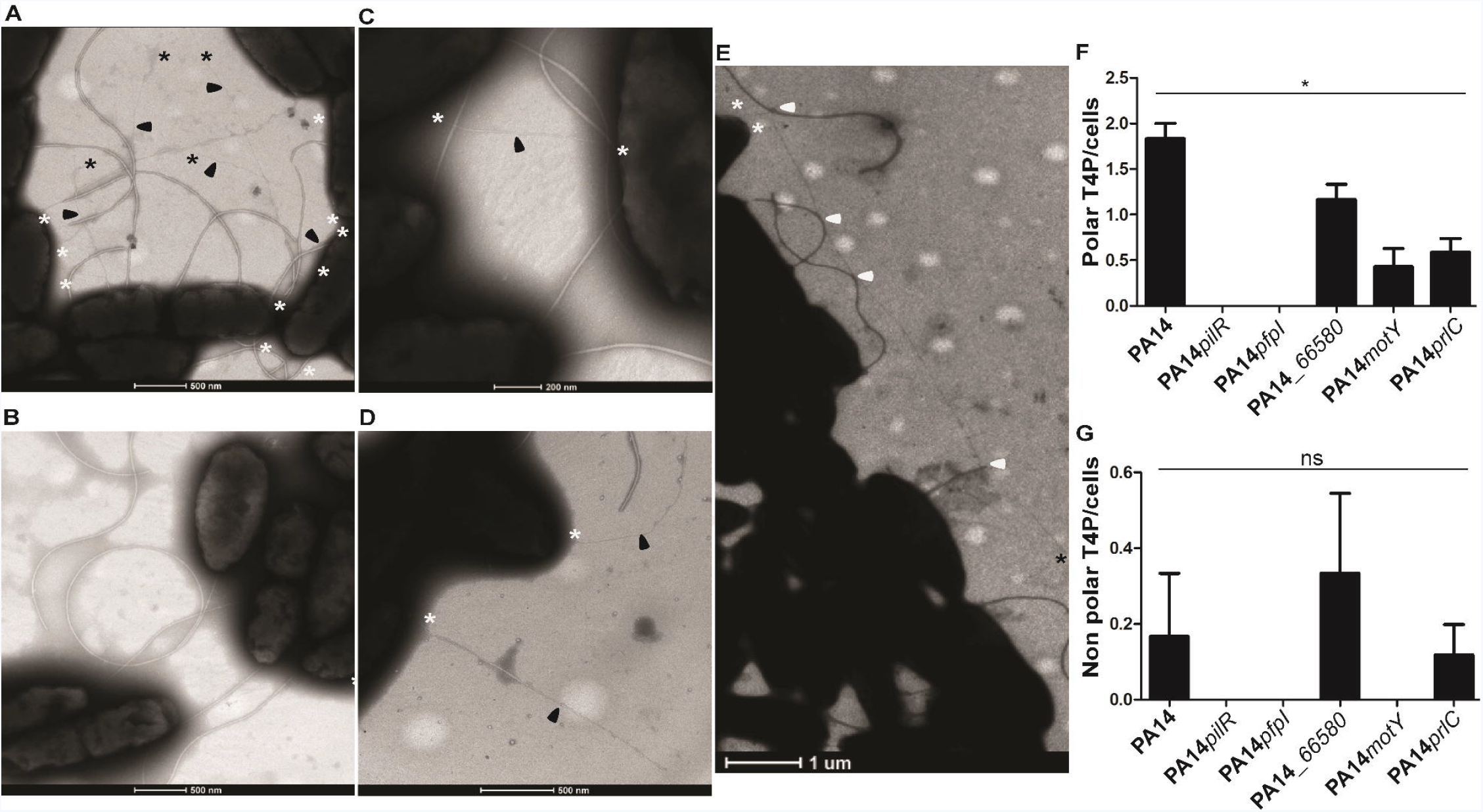
Visualizing T4P assembly and localisation. Representative images of (A) wildtype PA14, (B) a pili mutant *pilR*, (C) a *PA14_66580* mutant, (D) a *motY* mutant, and (E) a *prlC* mutant are shown; (F-G) quantification of T4P at the polar or non-polar cellular region/cells. All pili visualized and included in our analyses were between 1-2 μm in length. *pfpI* cells were indistinguishable from *pilR* as represented in (B); a *prlC* mutant had longer T4P compared to wildtype and in some cases these pili appeared to interact with the flagella (potential interactions marked with white arrows) (E). In each image the pili are arrowed in black, with a black asterix at any free ends and a white asterix where the pili appear to join or go under a cell membrane. Images are representative of triplicate grids imaged in biological triplicate. For each replicate of each strain at least 200 cells were visualised. In (F) Two-tailed student’s t-test, * (p<0.05) compared to wildtype; in (G) Two-tailed student’s t-test, ns determined for all samples compared to wildtype: vs. PA14*pilR* p = 0.363, vs. PA14*pfpI* p = 0.361, *PA14_66580* p = 0.611, PA14*motY* p = 0.363, PA14*prlC* p = 1.000.

### Investigating the role of *PA14_66580* in T4P assembly and function

A clean deletion in *PA14_66580* was generated in the orthologous gene (*PAK_05353* with the gene product having 99.82 % amino acid identity to PA14_66580) in the *P. aeruginosa* strain PAK. As was observed for the transposon mutant of *PA14_66580* in PA14, a reduction in twitching motility was also observed in the PAK deletion mutant PAK*05353* (Figure 3A). *PA14_66580/PAK_05353* is encoded just upstream of the *pilMNOP* gene cluster which encodes the components in the alignment subcomplex, and the outer membrane associated secretin complex of PilP and PilQ which is involved in T4P outer membrane extrusion (11-14). Additionally, PA14_66580/PAK_05353 is also annotated as a predicted ExeA-like protein. ExeA is an ATPase which binds peptidoglycan and is involved in transport and multimerization of ExeD into the outer membrane to form the functional secretin of the Type II secretion system (56). Given this, we hypothesized that PA14_66580/PAK_05353 may be involved in multimerization and/or localization of the PilQ secretin complex. To investigate this we performed immunoblotting of whole cell lysates of wildtype, PAK*05353* and PAK*pilQ* strains harvested from agar plates for both the multimeric and monomeric forms of PilQ (Figure 3B). This revealed that PAK*05353* was able to form both multimers and monomers of PilQ to the same extent as wildtype indicating that PAK05353 does not appear to play a role in PilQ multimerization.

**Figure 3.**
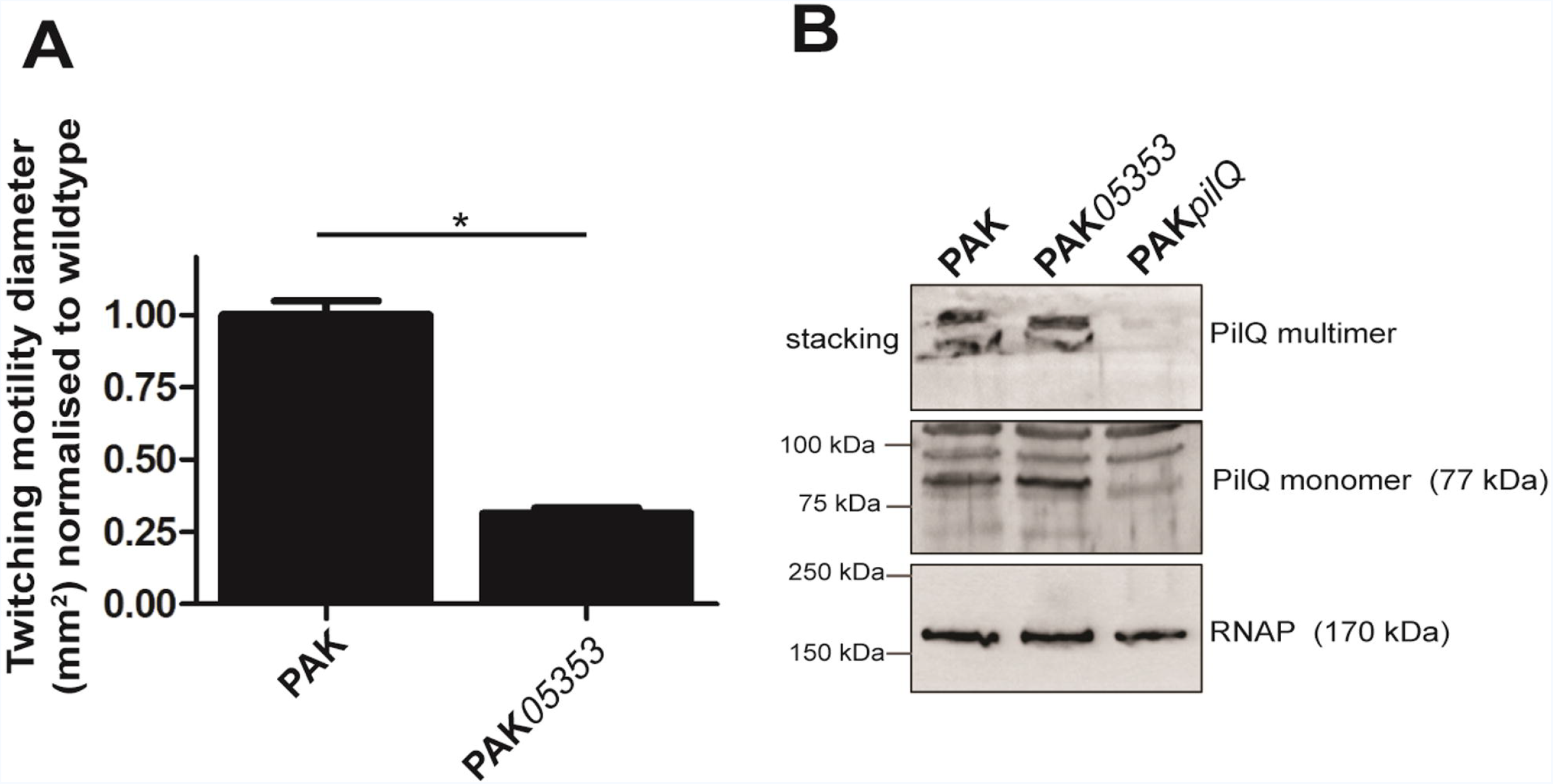
Twitching motility and PilQ secretin phenotypes in PAK*05353* (*PA14_66580* mutant). (A) Subsurface twitching motility-mediated interstitial biofilm expansion at agar/plastic interface after 48 h incubation at 37 °C for PAK and clean deletion of *PA14_66580* in PAK (PAK*05353*) is presented as the mean surface area in mm^2^ ± standard error of the mean normalized against wildtype as obtained from 3 independent experiments performed in triplicate. Two-tailed student’s t-test, * (p<0.0005) compared to wildtype. (B) Immunoblot of PilQ from whole cell preparations of strains PAK, PAK*05353* and PAK*pilQ* obtained from overnight (20 h) confluent lawns grown at 37 °C on LB agar plates. RNAP was used as a loading control.

### Functional Gene Enrichment Analysis

Enrichment analyses of genes that had increased or decreased mutant populations in the TraDIS output using the KEGG database (57) revealed that 3 key pathways were significantly altered. These were: flagella biosynthesis, two-component systems (TCS) and chemotaxis (Table S3 (increased population) and Table S4 (decreased population)).

We noticed that a number of flagella-associated structural and regulatory genes had altered mutant abundances following selection for twitching motility-mediated biofilm formation in our TraDIS assay, and some single mutants were confirmed to have significantly altered levels of twitching motility compared to wildtype (Figure 1A). Remarkably, this revealed a strong correlation between gene products predicted to have a negative effect on twitching motility (which corresponds to a positive log-fold change in our TraDIS output (Table S1), or a measured increase in twitching motility of the transposon mutant (Figure S2)) with proteins associated with the outer part of the cell envelope and thus the outer part of the flagellum body. In contrast, proteins associated with the inner part of the cell envelope and flagellum body were predicted to have a positive effect on twitching motility (which corresponds to a negative log-fold change in our TraDIS output (Table S1) or a measured decrease in twitching motility of the transposon mutant (Figure S2)) (Figure 4).

**Figure 4.**
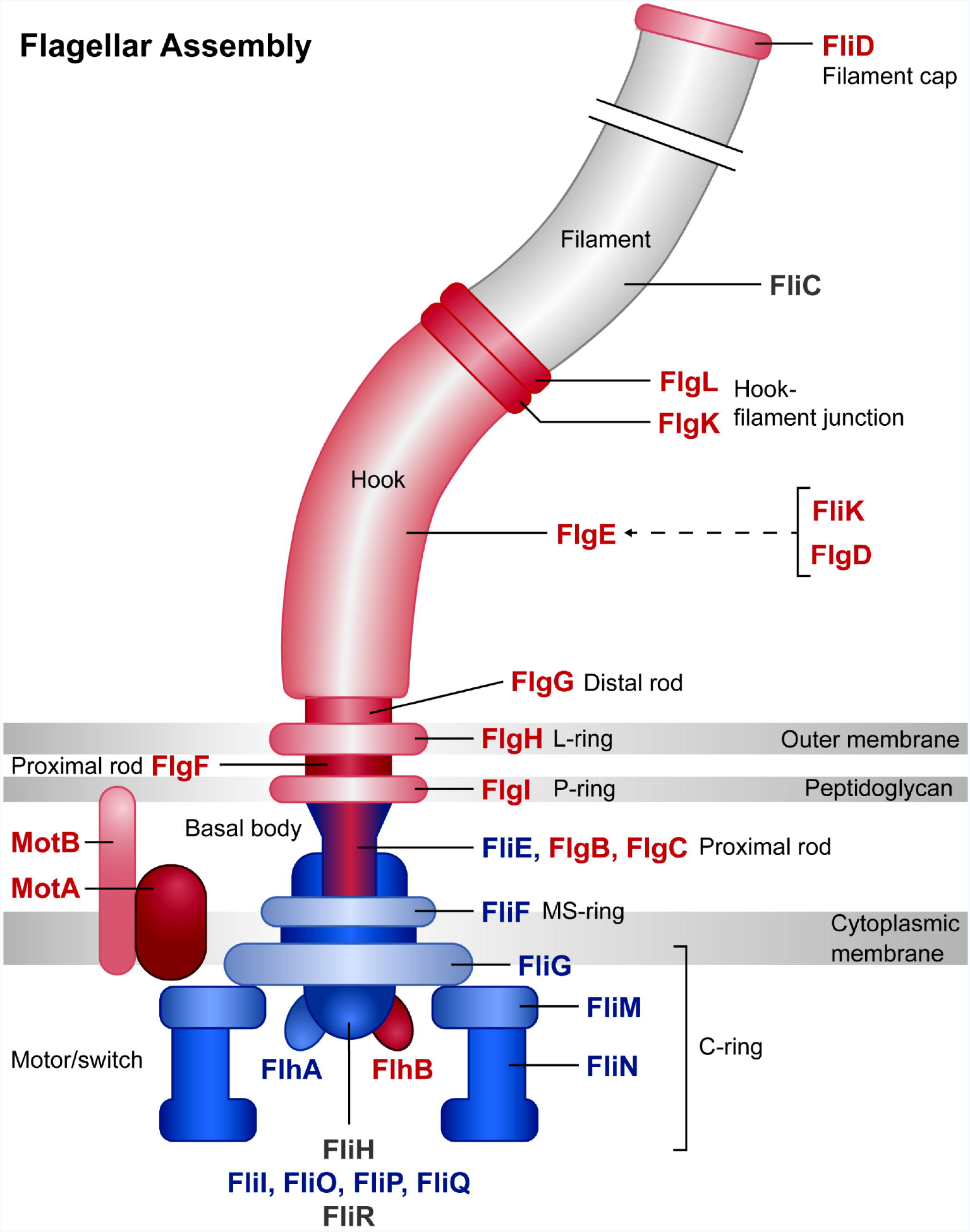
Relative log-fold change of transposon insertions in genes for flagella components. Gene products involved in flagella structure or regulation of flagella function are represented in this diagram. Genes which had a positive log fold change (which implies a negative effect on twitching motility) are coloured red, and are mostly located in the outer part of the cell envelope and flagellum body. Genes which had a negative log fold change (which implies a positive effect on twitching motility) are coloured blue and are mostly located in the inner part of the cell envelope and flagellum body. Output image generated from KEGG Mapper tool (http://www.kegg.jp/kegg/mapper.html).

The chemotaxis pathway identified in our functional gene enrichment analysis included mutants of swimming chemotaxis (*che*) genes which appeared to promote (*cheA/B/Z/Y*) as well as inhibit (*cheR/W*) twitching motility-mediated biofilm formation. This suggests a balance between bacterial chemotaxis and twitching motility, especially as the chemotaxis pathway also controls flagella assembly. The TCS linked to twitching motility were mostly known genes, for instance *algZ/R* involved in alginate biosynthesis, or the *pil* genes in T4P production, but also included some unexpected genes related to osmotic stability, such as *cusS/R* involved in copper efflux, or *dctA/B/D/P* for C4-dicarboxtrate transport.

## Discussion

In this study, we have successfully applied a physical separation-based TraDIS approach to identify genes involved in twitching motility-mediated biofilm formation in *P. aeruginosa*. Using this method, we could detect almost all genes currently known to be involved in T4P assembly and twitching motility, in addition to a large number of genes identified in our TraDIS output (Table S1) and a select group for further study (Table 2) not previously known to be involved.

A functional enrichment analysis of all genes that have altered mutant abundances in our assay identified 3 major groups of gene function that were affected during twitching motility: flagella assembly, bacterial chemotaxis and TCS. Perhaps the most interesting from these is the potential involvement of the flagella as suggested from the predicted (Table S1) or determined (Figure S2) differential effect of structural and regulatory flagella components on twitching motility. Specifically, we observed a strong correlation between gene products predicted to have a negative effect on twitching motility with proteins associated with the outer part of the flagella body, and in contrast, proteins associated with the inner part of the flagella body were predicted to have a positive effect on twitching motility (Figure 4). This is intriguing as it suggests a differential effect on twitching motility by flagella components based upon their cellular location and certainly warrants further investigation in future work.

For each of the 11 gene targets selected for further investigation (Table 2) the twitching motility phenotype was confirmed in a sub-surface stab assay, with growth assays performed to demonstrate that the observed twitching phenotype was not due to a growth defect. Submerged biofilm formation was also assayed which revealed that only *kinB* had a significant decrease in levels compared to wildtype (Figure 1B), demonstrating that our TraDIS assay did indeed selectively identify genes specific for twitching motility-mediated biofilm expansion on a semi-solid surface. Overall these assays confirmed the twitching motility phenotype observed for *prlC*, *PA14_66580, pfpI, fliG* and *motY* was not due simply to a growth related defect. For these mutants TEM was used to determine whether the twitching motility phenotype could be attributed to alterations in levels and/or localization of surface assembled T4P. No pili were observed in a *pfpI* mutant (Figure 2B), which explains the observed lack of twitching motility (Figure 1A). PfpI is an intracellular protease which affects antibiotic resistance, swarming motility and biofilm formation in *P. aeruginosa* (58) however, to our knowledge the current study is the first report to a role for PfpI in twitching motility. Given the established role of intracellular proteases in controlling levels of a range of chaperones and regulatory proteins it is likely that the protease activity of PfpI is required for control of regulators or other proteins involved in T4P biogenesis and/or assembly.

A role for PA14_66580 was also investigated in the formation of the PilQ secretin to allow T4P extrusion and thus function. This was based upon the proximity of *PA14_66580* to the *pilMNOP* operon, which encodes components that link the outer membrane PilQ secretin to the inner membrane motor complex. Additionally PA14_66580 possesses the same conserved domain as ExeA (Uniprot: http://www.uniprot.org/uniprot/A0A0H2ZID1), which is an ATPase that binds peptidoglycan and is involved in transport and multimerization of ExeD into the outer membrane to form the secretin of the Type II secretion system (56). Our TEM data revealed that *PA14_66580* had reduced numbers of pili compared to wildtype (Figure 2C, F), which correlates with the observed reduction in twitching motility in both PA14 and PAK strain backgrounds (Figure 1A and Figure 3A). Given that no difference in the expression of monomeric or multimeric PilQ was observed in a mutant of *PA14_566580* in PAK (PAK*05353*) (Figure 3B), we suggest that the reduction in surface T4P and twitching motility levels is not due to a lack of secretin formation. PA14_66580 could instead be involved in stabilization of the secretin pore and/or formation of the assembly and motor subcomplexes in order to allow full functionality of the T4P.

A *prlC* mutant was found to have increased levels of twitching motility compared to wildtype (Figure 1A), a reduction in polar surface assembled T4P (Figure 2F) and to have a putative interaction between the surface-assembled flagella and T4P (Figure 2D). PrlC is uncharacterized in *P. aeruginosa* however it has an M3 peptidase domain (Pfam PF01432) which is associated with mammalian and bacterial oligopeptidases. The homologue in *E. coli* is a cytoplasmic protease (also named PrlC) which appears to be a partner in degradation of peptides produced by ATP-dependent proteases from multiple protein degradation pathways (59). A homologue of PrlC also exists in *Aeromonas hydrophilia*. A mutant of the oligopeptidase *pepF* was shown to have decreased swimming motility, increased biofilm formation in a crystal violet microtitre plate assay and increased attachment to epithelial cells (60). While the exact role of PepF has been not elucidated, this published data suggests that PepF could be involved in processing proteins involved in biogenesis or regulation of the polar flagella, used for swimming, or the bundle-forming pili (Bfp) or Type-IV *Aeromonas* pili (Tap) used for attachment. While it is unclear exactly how PrlC in *P. aeruginosa* is influencing twitching motility, our results suggest a similar role as for PepF in *A. hydrophilia* in processing proteins involved in biogenesis, assembly or regulation of the T4P. Alternatively, given the putative interaction of the flagella and T4P observed (Figure 2D), PrlC may be involved in processing flagella-associated proteins to ultimately affect the putative interaction between these two motility machines and thus the function of the T4P (as suggested from Figure 4).

We observed that a *motY* mutant had increased twitching motility levels (Figure 1A) but reduced levels of T4P (Figure 2A, F). MotY is a peptidoglycan binding protein which is required for MotAB-mediated flagella motor rotation and is associated with the outer-membrane (61). While a *motY* mutant is severely impaired for flagella-mediated motility on semi-solid surfaces (swarming motility), there is only a slight decrease in levels of swimming motility compared to wildtype in liquid media (61). The *motY* mutant used in this study (47) has a transposon insertion in the centre of the protein, and thus lacks the C-terminus of the protein, which has been shown to be involved in stabilizing the association of MotY with the stator proteins MotAB to allow flagella rotation (62). Given the observed reduction in surface assembled pili (Figure 2A, F) and increase in twitching motility (Figure 1A) this suggests that, as discussed above, a defect in flagella function is likely affecting the function of the T4P to produce the observed twitching motility phenotype.

FliG is one of the proteins in the rotor-mounted switch complex (C ring), located at the base of the basal body in the cytoplasm, and is important for directing flagella rotation (30). We observed a decrease in twitching motility of *fliG* (Figure 1A). Deletion of the C-terminal region of FliG (as is the case for our *fliG* transposon mutant (47)) results in a strain which produces non-functional flagella (63). Thus we would predict that, as for *motY*, our *fliG* mutant would also have wildtype levels of assembled T4P, with the presence of non-functional flagella in *fliG* also potentially affecting function of the T4P in twitching motility.

This study has identified both known and novel components involved in twitching motility in *P. aeruginosa*. Additionally, we have provided analyses which suggest a differential effect of flagella proteins on T4P function based upon their cellular location and points towards a possible interaction between the flagella and T4P machines to influence twitching motility. Overall these results highlight the success of our TraDIS-based approach and point to a number of intriguing new players involved in twitching motility-mediated biofilm expansion in *P. aeruginosa*.

## Acknowledgements

The Authors state that they have no conflict of interest.

L.M.N. is supported by MRC Grant MR/N023250/1 and a Marie Curie Fellowship (PIIF-GA-2013-625318). A.F. is supported by Medical Research Council (MRC) Grants MR/K001930/1 and MR/N023250/1 and Biotechnology and Biological Sciences Research Council (BBSRC) Grant BB/N002539/1. A.K.C and C.J.B were supported by the Medical Research Council (Grant G1100100/1). This work was supported by an MRC Centenary Award (Grant G1100189) and the Wellcome Trust (Grant WT098051).

The authors are grateful to Stephen Lory for gifting the clean deletion mutant of PAKΔ*pilQ* and the α-PilQ antibody and to Sandy Pernitzsch / Scigraphix for assistance with Figure 4.

## Supplementary Table Legends

**Table S1. Full list of all gene hits obtained in TraDIS screen.** Table contains PA14 gene locus tag, gene name, annotated gene function, the logFC (log2 fold change) in transposon insertions from mutants obtained from the inner (non-twitching) zone vs mutants obtained from the outer (active-twitching) motility zone, and the statistical q value for all replicates.

**Table S2. Full list of PA14 transposon mutants used in the current study.** PA14 mutants from the PA14 transposon mutant collection (47) with gene name, annotated gene function and the plate and well where the mutant was taken from in the collection.

**Table S3. Enrichment analyses of genes that had an increased mutant population in the TraDIS output.** KEGG database (57) was used to identify pathways enriched in mutants with increased mutant populations in the TraDIS output (Table S1).

**Table S4. Enrichment analyses of genes that had a decreased mutant population in the TraDIS output.** KEGG database (57) was used to identify pathways enriched in mutants with decreased mutant populations in the TraDIS output (Table S1).

### Supplementary Figure Legends

**Figure S1.**
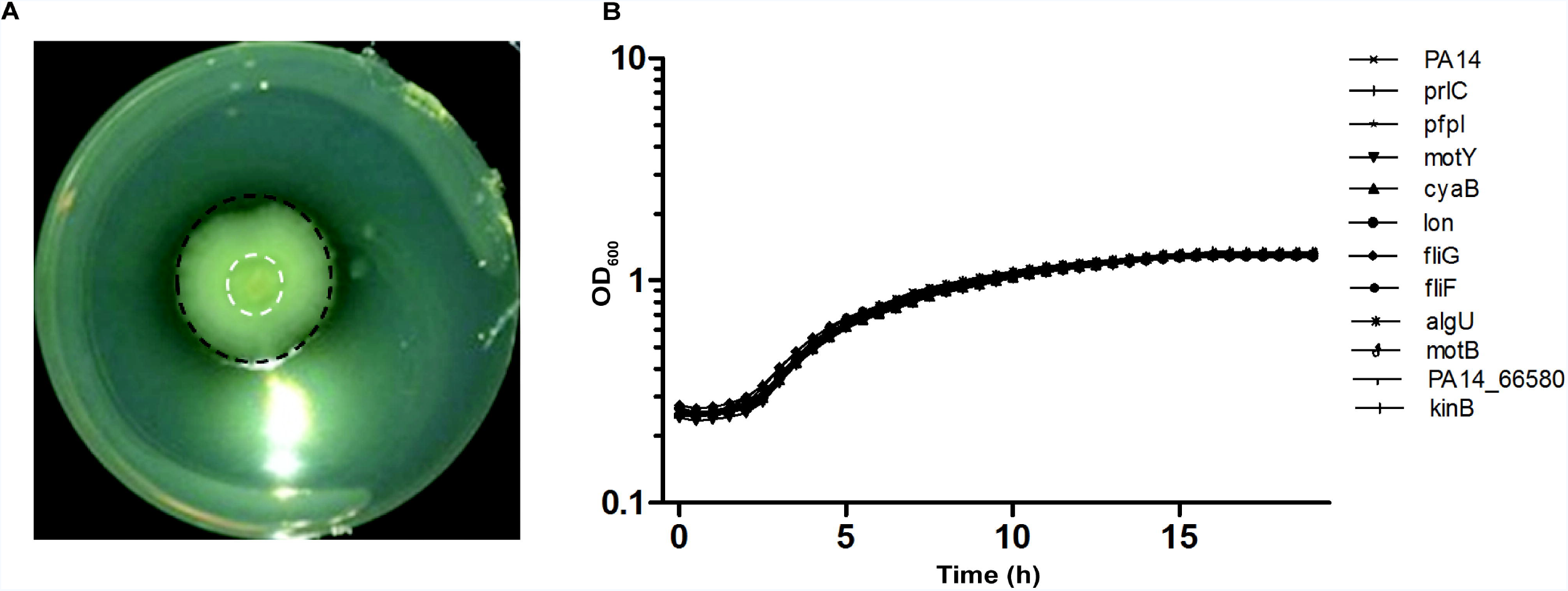
(A) The transposon mutant pool was inoculated in the centre of an inverted agar plate and incubated for 72 h at 37 °C. Mutants were separated based upon their ability to undergo twitching motility-mediated biofilm expansion away from the inoculation site. After 72 hrs mutants were harvested from the inner non-motile region (outlined in white) and the outer active twitching edge (outlined in black). Genomic DNA was extracted from each pool of mutants and sequenced using a mass parallel approach. (B) Growth rates in minimal media for 11 selected transposon mutants assayed for a submerged biofilm defect in Figure 1B. Growth rates were determined by incubation of transposon mutants at 37 °C for 19 h in M63 minimal media (same media used for submerged biofilm assays). There was no significant different between growth rates of transposon mutants compared to wildtype predicted by a one-way ANOVA with Dunnett’s multiple comparison test. Data are represented as the mean ± standard deviation for 2 independent replicates performed in triplicate.

**Figure S2.**
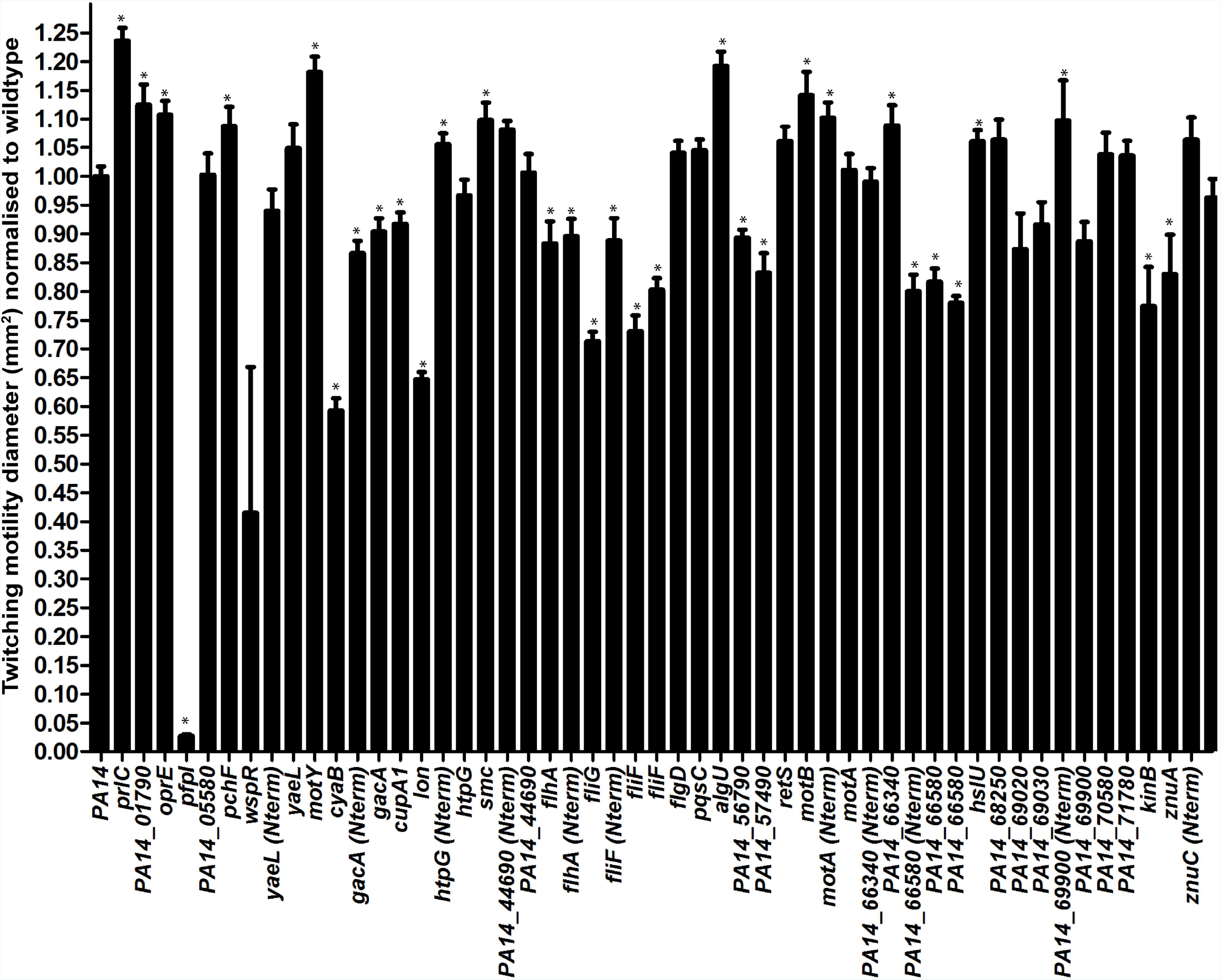
Twitching motility of selected transposon mutant targets identified using TraDIS. Sub-surface twitching motility-mediated interstitial biofilm expansion at agar/plastic interface after 24 h incubation at 37 °C is presented as the average surface area in mm^2^ ± standard deviation normalized against wildtype as obtained from 2 independent experiments performed in triplicate.

